# The coordination of hip, knee and ankle joint angles during gait in soccer players and controls

**DOI:** 10.1101/2021.09.24.461658

**Authors:** Morteza Yaserifar, Ziya Fallah Mohammadi, Sayed Esmaeil Hosseininejad, Iman Esmaili Paeen Afrakoti, Kenneth Meijer, Tjeerd W Boonstra

## Abstract

**Background:** Clinical researchers are trying to unravel the impact of different training interventions on the kinematics of human gait. However, the effects of long-term training experience on the kinematics of a healthy gait pattern remains unclear. Here we assess the effect of long-term training experience on joint angle variability during walking.

**Methods:** Hip, knee, and ankle joint angles from fourteen soccer players and sixteen controls were acquired during treadmill and overground walking. Hip-knee coupling, knee-ankle coupling and coupling angle variability (CAV) of the right leg in the sagittal plane were assessed using a vector coding technique.

**Results:** Soccer players showed reduced hip-knee CAV during the mid-stance and terminal-stance phases and reduced knee-ankle CAV during the pre-swing phase of gait compared to the control group. In addition, soccer players less often used an ankle coordination pattern, in which only the ankle joint but not the knee joint rotates.

**Interpretation:** These findings show that soccer players had more stability in the ankle joint during the stance phase of the gait compared to the control group. Future studies can test whether these differences in the coordination of the ankle joint reflect the effects of long-term training on normal gait by comparing knee-ankle coupling and variability before and after exercise training interventions.

## 1. Introduction

It is well documented that exercise training improves cardiovascular function, stability, and strength (Sherrington et al., 2008). These training effects are particularly pronounced in older people or people with disabilities. The advantage of exercise training in improving the functional capacity of older adults have been recently demonstrated (Font-Jutglà, Gimeno, Roig, da Silva, & Villarroel, 2019; Xia et al., 2020). For instance, in multi-component exercise program, resistance training contributes most to overall enhancements of muscle strength during acute hospitalization (Sáez de Asteasu et al., 2020). Furthermore, exercise training has an important effect on movement rehabilitation in patients and can help them to resume their routine life. For example, virtual reality training could help to extend the balance and walking abilities of patients (Bang, Son, & Kim, 2016). Enhancement of daily life activity was one of the most important outcomes of this study and walking is a key activity that can help all groups achieve an independent life.

Human walking pattern is unique compared to primate gait (Maurice Abitbol, 1988; Rodman & McHenry, 1980). To enable this unique ability some adaptations have taken place in the human body: during a normal walking, the head and center of gravity are lowest near toe-off and highest at mid-stance (Bramble & Lieberman, 2004), so that we can walk smoothly and highly efficiently for a long time. As human walking is a complex motion, it requires coordination between multiple body segments: Multiple lower limb joints and segments should move synchronously during normal gait (DeLeo, Dierks, Ferber, & Davis, 2004), and disruptions in coordination, either internal or external, may result in impaired gait. For example, weakness of one or a group of muscles can create an abnormal pattern and change the kinematics of gait (Galli et al., 2012). Likewise, trial-to-trial variability in movement pattern are associated with skill level or expertise (Wilson, Simpson, Van Emmerik, & Hamill, 2008), and optimal variability in movement patterns is a characteristics of healthy functioning (Harbourne & Stergiou, 2009).

Gait variability can be quantified at several levels, such as kinematics and stride characteristics (Ulman, Ranganathan, Queen, & Srinivasan, 2019). The relative motion between the angular time series of two joints has been used to distinguish normal from disordered or symmetrical from asymmetrical gait patterns and has also been applied to assess how coordination in sports differs as a function of expertise (Glazier, 2006). It has been shown that coordination variability decrease when skilled athletes perform more consistently or better regulated (Wilson et al., 2008). Vector coding is a non-linear technique to quantify coordination and variability. It quantifies the continuous dynamic interaction between segments by detecting the vector orientation of the angle-angle diagram relative to the horizontal (Hamill, Haddad, & McDermott, 2000; Sparrow, Donovan, Van Emmerik, & Barry, 1987). It is used to estimate the coupling angle variability (CAV) between body segments and the CAV can reveal changes in coordinative state between different groups (Heiderscheit, Hamill, & van Emmerik, 2002).

Although the functional role of short- and long-term motor learning on movement variability has been investigated (Bartlett, Wheat, & Robins, 2007; Bradshaw, Maulder, & Keogh, 2007; Wu, Miyamoto, Castro, Ölveczky, & Smith, 2014), the effects of long-term training on the joint variability of the lower extremities during gait remain unclear. While training programs could differ between sports, walking often constitutes a considerable part of training programs, in particular for soccer players (Krustrup, Mohr, Ellingsgaard, & Bangsbo, 2005; Rampinini, Coutts, Castagna, Sassi, & Impellizzeri, 2007). Understanding the effects of long-term training on gait is important when applying rehabilitation training in different groups with and without exercise training experience.

In this study, we used the vector coding technique and CAV to assess differences in coordination and variability of right leg joints in the sagittal plane between soccer players and controls. Long-term soccer training may change the control of general movements such as gait. As gait is an important part of soccer training program, we expect that soccer players show reduced lower extremity joint variability when they walk on a treadmill or overground compared to the control group. Being able to distinguish differences in joint kinematics between soccer players and controls may contribute to our understanding of the effects of long-term training on movement coordination.

## 2. Methods

### 2.1. Participants

Thirty male participants, fourteen soccer players (height: 175 ± 4 cm, mass: 70.2 ± 8.0 kg, age: 23 ± 5 years) and sixteen non-athletes (height: 175 ± 7 cm, mass: 79.2 ± 16.0 kg, age 24.7 ± 5 years) with no history of musculoskeletal injury gave written agreement to participate in the study. The athletes had at least seven years continuous soccer training experience and were members of the soccer team of University of Mazandaran. The control group reported that they did not have a history of doing regular exercise training or injuries that could affect their walking. All participants were students at the University of Mazandaran. The study protocol was approved by the Office of Research Ethics at the University of Mazandaran and prior to beginning the protocol, participants provided written informed consent.

### 2.2. Experimental setup and procedure

Participants wore comfortable walking shoes and performed treadmill walking (H/P COSMOS treadmill, Germany) and overground walking on a 10-meter walkway. Their preferred walking speed was recorded during the familiarization phase. Overground walking speed was recorded using a stopwatch and determined from the mean of three recordings while walking on the 10 m walkway. Treadmill walking speed was directly recorded by a monitor on the treadmill when a participant indicated they were walking at their preferred speed. All participants first completed overground walking and then treadmill walking (see Fig. 1).

**Fig. 1.**
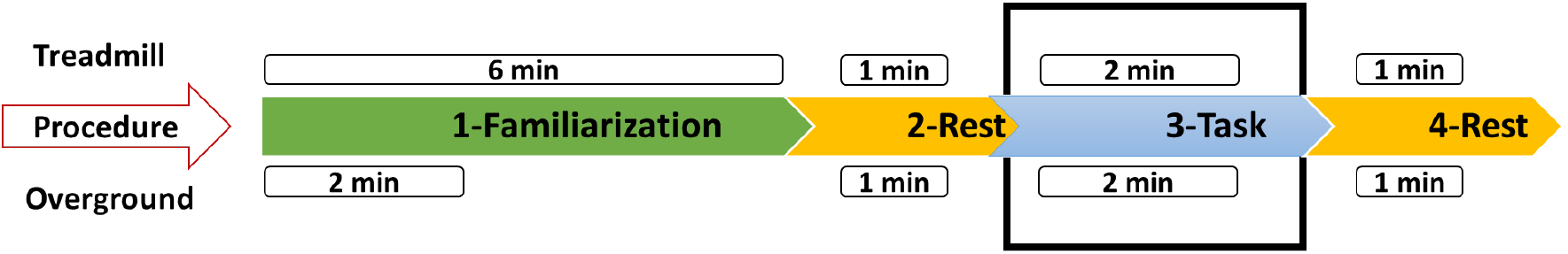
Schematic of treadmill and overground walking procedure. The procedure was divided into four phases: familiarization, rest, task, and rest phases, respectively. The black box line shows when data was collected. The time duration of treadmill and overground walking during each step are distinguished on the top and down of the procedures, respectively. During rests participants were sitting on a chair.

### 2.3. Data acquisition

Kinematic data were recorded using a 3D inertial measurement unit (IMU) consisting of magnetometers, accelerometers and gyroscopes (Noraxon MyoMotion system, USA). We used five sensors to collect 3-dimensional kinematics data during walking: three sensors on thigh, shank, and foot of right leg and one sensor on lumbar spine to capture hip, knee, and ankle joint angles and foot switch data during walking (Mundt et al., 2019). A sensor was placed on lower thoracic (T12) to record lower spinal horizontal rotation during overground walking. All sensors were placed based on Noraxon MyoMotion setup and sampled at 100 Hz.

### 2.4. Data analysis

Three-dimensional hip, knee, and ankle angles were processed using a low-pass Butterworth filter with a cut-off frequency of 6 Hz (Winter, Sidwall, & Hobson, 1974). Participants had to turn at the end of the 10-meter walkway and continued walking for two minutes. To extract the data from the straight walkway, excluding turning points, we applied the method that we used previously (Yaserifar et al., 2021). Briefly, we collected the lower spinal horizontal rotation data to remove data during turning points and only analyzed data during straight line walking: The zero-crossings indicated the middle of the turning movement and we removed 1.5 s before and after each zero-crossing. The remaining data was used for further analysis. The gait cycles of right dominant leg were segmented using footswitch data for both overground and treadmill walking. Each cycle was defined by consecutive heel strikes and was time normalized and scaled to 100% of the gait cycle.

### 2.5. Calculation of coupling angle and coupling angle variability

Custom MATLAB (Version 2018A, USA) scripts were developed to estimate the coupling angles and CAV. The coupling angles and CAV were assessed in sagittal plane (Luc-Harkey et al., 2016). Reliable estimation of individuals’ variability requires at least ten stride cycles (Hafer & Boyer, 2017), and we extracted data from the joint angles of fifty gait cycles during treadmill and overground walking. Angle-angle diagram were then created with the proximal joint on the horizontal axis and the distal joint on the vertical axis. Vector coding techniques was used to estimate the coupling angle and joint angle variability of fifty continuous gait cycles based on consecutive proximal and distal joint angles (Peters, Haddad, Heiderscheit, Van Emmerik, & Hamill, 2003). The coupling angle (*γ*_*i*_) was calculated according to equations (1) and (2) for each time point (i) within the gait cycle by the vector orientation between two adjacent data points in time on the angle–angle diagram relative to the right horizontal (Needham, Naemi, & Chockalingam, 2014).

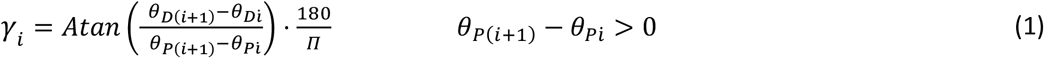

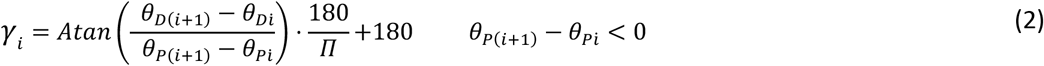

The following conditions (3) were applied:

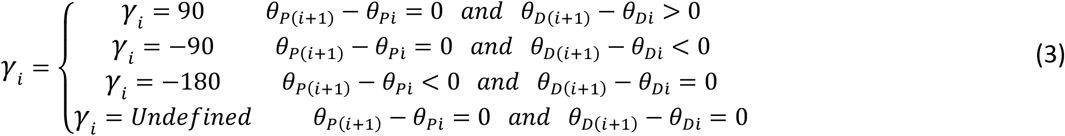

The coupling angle was then corrected to obtain a value between 0° and 360° (Chang, Van Emmerik, & Hamill, 2008; Sparrow et al., 1987):

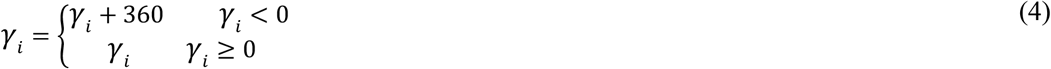

As these angles are obtained from a polar distribution, the mean coupling angles 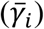 must be extracted using circular statistics. So, 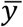 were calculated using the average of horizontal and vertical components (Batschelet, 1981; Hamill et al., 2000):

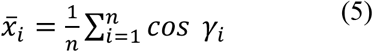

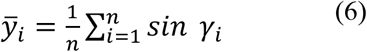

We then corrected the average coupling angle to obtain a value between 0° and 360° (7) (Needham et al., 2014).

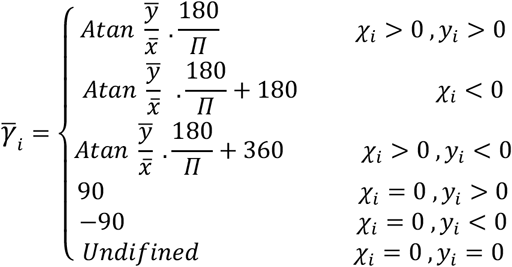

Mean 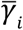 were then categorized into one of four coordination patterns: in-phase, anti-phase, proximal phase and distal phase (Chang et al., 2008). When the coupling angles are 45° and 225°, the coupling is in-phase, and both joints rotate in the same direction, whereas if two joints rotate in the opposite direction, i.e. at 135° and 315°, this is considered anti-phase coupling. When coupling angles parallel the horizontal 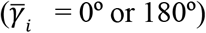, there is rotation of the proximal joint but not the distal joint and this is considered the proximal-phase pattern. Finally, vertically directed coupling angles 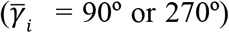 indicate a distal-phase pattern, in which only the distal joint rotates (Chang et al., 2008). We determined the frequency with which each coordination pattern occurred at different phases of the gait cycle.

The length of the mean vector was then calculated according equation (8), and the coupling angle variability (*CAV*_*i*_) was determined according to equation (9) (Batschelet, 1981).

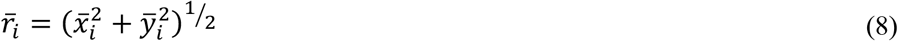

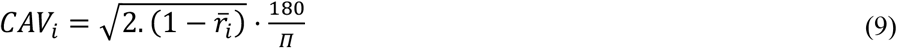

To assist in the interpretation, the gait cycle was divided into seven sub-phases (Eggleston, Harry, Hickman, & Dufek, 2017): loading response (1-10%), mid-stance (11-30%), terminal stance (31-50%), pre-swing (51-60%), initial swing (61-73%), mid-swing (74-87%), and terminal swing (88-100%). The percentages refer to the percentage of gait cycle. The mean in each sub-phase was used for statistical analysis.

### 2.6. Statistical analysis

A univariate analysis of covariance (UNIANCOVA) was used to compare the coupling angle, CAV, and coordination pattern frequency across groups and surfaces. In the model we included group (soccer players and control group) and surface (treadmill and overground) as fixed effect factors. As participants walked at their own preferred speed, walking speed was added as a covariate. The coupling angle, CAV, and coordination frequency were assessed at seven sub-phases of the gait cycle and we used the Benjamini-Hochberg procedure to adjust the p-values for multiple testing (Benjamini & Hochberg, 1995). The level of statistical significance was set at α = 0.05.

## 3. Results

Walking speed was higher in soccer players compared to controls and during overground walking compared to treadmill walking (Fig. 2), as reflected by the significant main effects of group (F_1,56_ = 10.7, P_adj_ = 0.03) and surface (F_1,56_ = 159.3, P_adj_ < 0.005). The interaction effect was not significant (P>0.9).

**Fig. 2.**
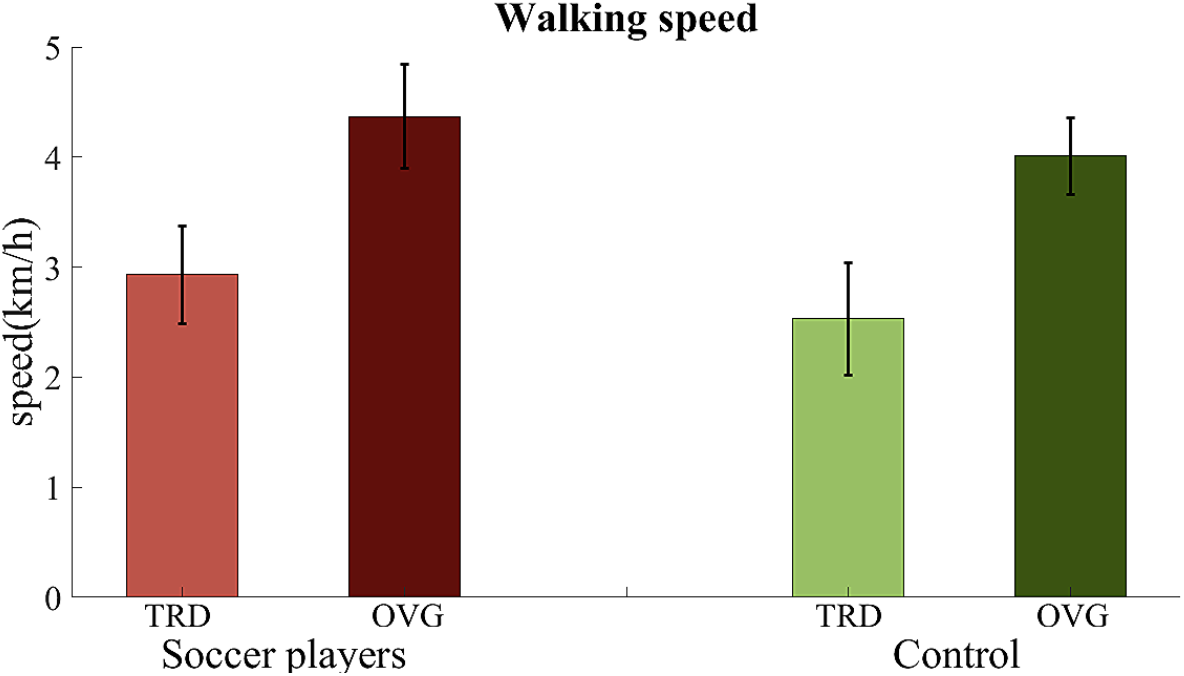
Walking speed of soccer players and control group during walking overground (OVG) or on a treadmill (TRD). Error bars show the standard deviation between participants.

Mean joint angles for both groups are shown in Fig. 3. Although the pattern is very similar in both groups and surfaces, the ankle angle reveals some differences between conditions (Fig. 3C). In particular, the pattern of angular variations in the ankle joint of soccer players on the treadmill in the propulsion and swing phases appears to show more plantarflexion.

**Fig. 3.**
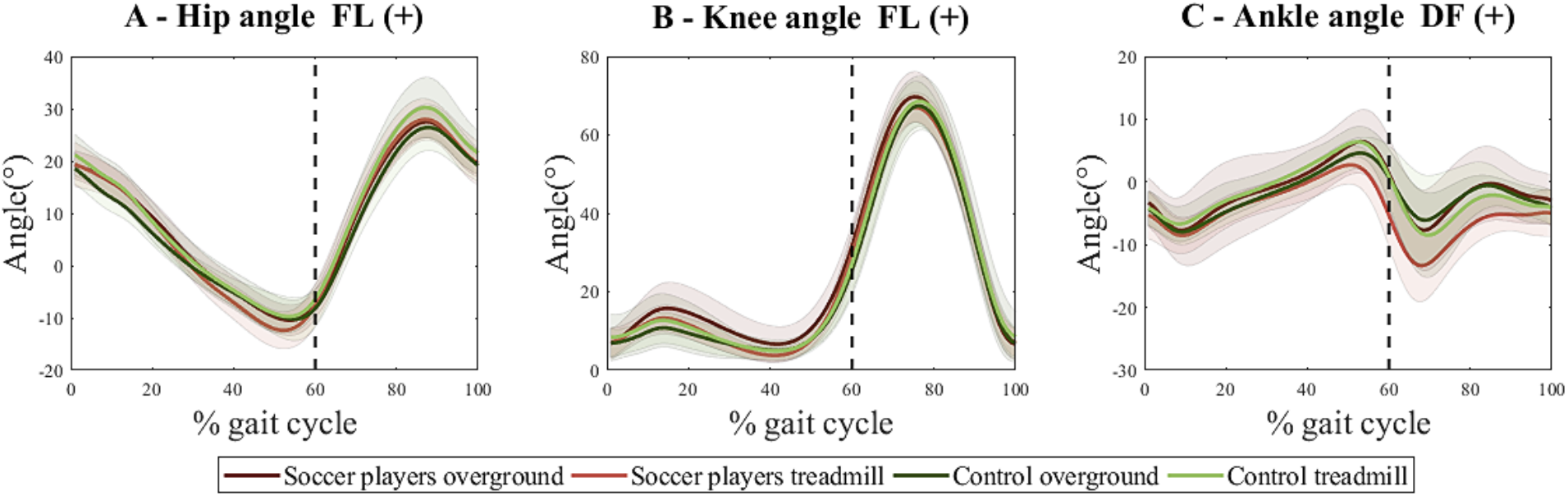
Joint kinematics data in sagittal plan of soccer players and controls during walking. Left panel shows the hip angle, middle panel shows the knee angle, and the right panel shows the ankle angle during a full gait cycle. The black dash line indicates toe-off and separates the stance phase (left side) and swing phase (right side). Dark red, light red, dark green and light green color show soccer players overground, soccer players treadmill, controls overground and controls treadmill, respectively.

Inter-segmental coordination was assessed based on the angle-angle plots of the hip-knee and knee-ankle pairs (Fig. 4A and 4E). The coupling angle and CAV were computed from the vector orientation between two adjacent data points in time in the angle-angle diagram. At the beginning of gait cycle the coupling angle of the hip-knee pair starts in anti-phase coordination where there was knee flexion and hip extension during loading response phase (with hip and knee angles around 20 and 5 degrees, respectively). The coordination pattern then changed to a proximal-phase (hip) pattern in the control group during the mid-stance and terminal stance phases, whereas soccer players kept an in-phase coordination pattern during these phases (Fig. 4B, sections b and c). Both groups showed nearly the same trajectory of hip-knee coupling for the following phases of the gait cycle finishing with in-phase coordination (Fig. 4B, sections e and g). The coupling angle variability slowly decreased during the gait cycle with short increases mid-swing and before heel strike (Fig. 4C). Both groups showed similar coordination patterns during the gait cycle (Fig. 4D).

**Fig. 4.**
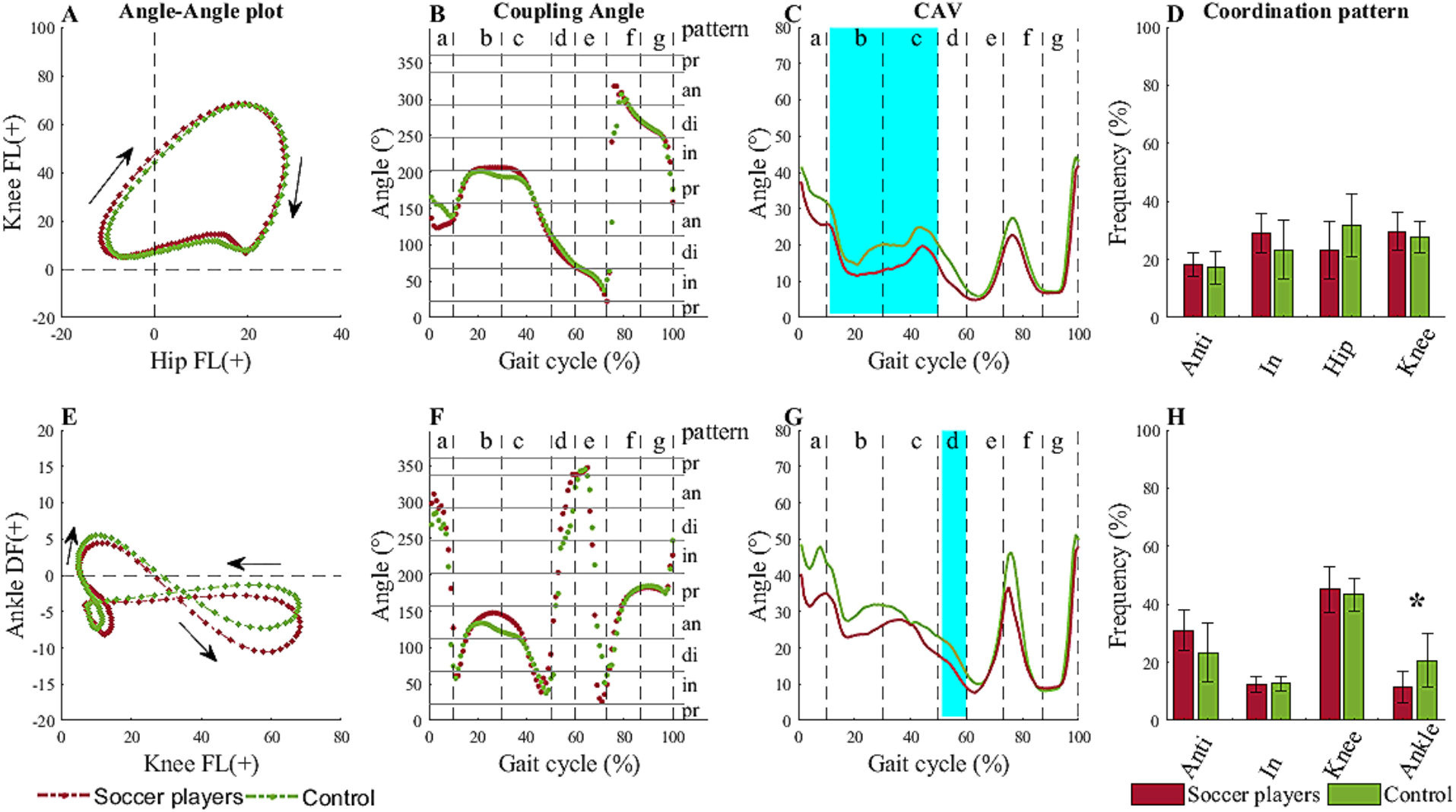
Hip-knee and knee-ankle coordination in sagittal plan for soccer players and controls. First column from left shows the angle-angle plots for hip-knee and knee-ankle coordination with proximal joint on the horizontal axis. The second column shows hip-knee and knee-ankle coupling angles. Note that the vector coding procedure yields coupling angle values between 0° - 360°. Vertical dash lines divide the gait cycle in seven sub-phases and horizontal grey lines show the associated coordination patterns (anti-phase, in-phase, proximal segment phase, distal segment phase). The third column shows coupling angle variability which is divided in gait cycle sub-phases (vertical dash lines). The cyan bands show the sub-phase with a significant main effect on CAV. The fourth column shows the average frequency (%) of each coordination patterns and the error bars show the standard deviation between participants. Anti = anti-phase, In = in-phase, a = loading response, b = mid-stance, c = terminal stance, d = pre-swing, e = initial swing, f = mid-swing, g = terminal swing. * = significant difference (P<0.05) in coordination pattern.

The knee-ankle coupling revealed a different trajectory during the gait cycle (Fig. 4E): There was a knee flexion versus ankle plantar flexion at the beginning of gait cycle (with a knee and ankle angles around 5 and −4 degrees, respectively). Soccer players initiated the gait cycle with an anti-phase coordination pattern 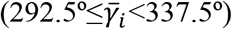 whereas controls initiated it with a distal-phase (ankle) pattern (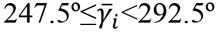; Fig. 4F, a). There was a different coupling pattern across the second part of mid-stance and the first part of terminal stance, but both groups shared a similar anti-phase coordination pattern (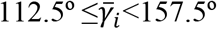; Fig. 4F, b and c). Furthermore, soccer players showed more scattered knee-ankle coupling during pre-swing phase 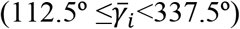 (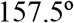 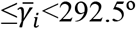; Fig. 4F, d) and less ankle coordination pattern compared to controls (Fig. 4H).

Figure 5 shows the mean coupling angles and CAV of soccer players and controls during the different phases of the gait cycle. When comparing the CAV both groups, soccer players showed decreased hip-knee CAV during the mid-stance and terminal stance phases (F_1,55_ =14.7, P_adj_ = 0.008 and F_1,55_ =18.2, P_adj_ = 0.003, respectively), and decreased knee-ankle CAV during the pre-swing phase (F_1,55_ =12.1, P_adj_ = 0.02) compared to controls (Fig. 4C, sections b and c, and Fig. 4G, section d). Furthermore, the knee-ankle coupling in soccer players revealed less often a distal-phase (ankle) ankle coordination pattern (F_1,55_ =14.4, P_adj_ = 0.008) during the gait cycle than controls (Fig. 3F). There were no significant differences for other sub-phases between groups or between surfaces. All statistical results of coupling and CAV and descriptive and statistical results of coordination pattern frequency are presented in Tables S1, S2 and S3, respectively.

**Fig. 5.**
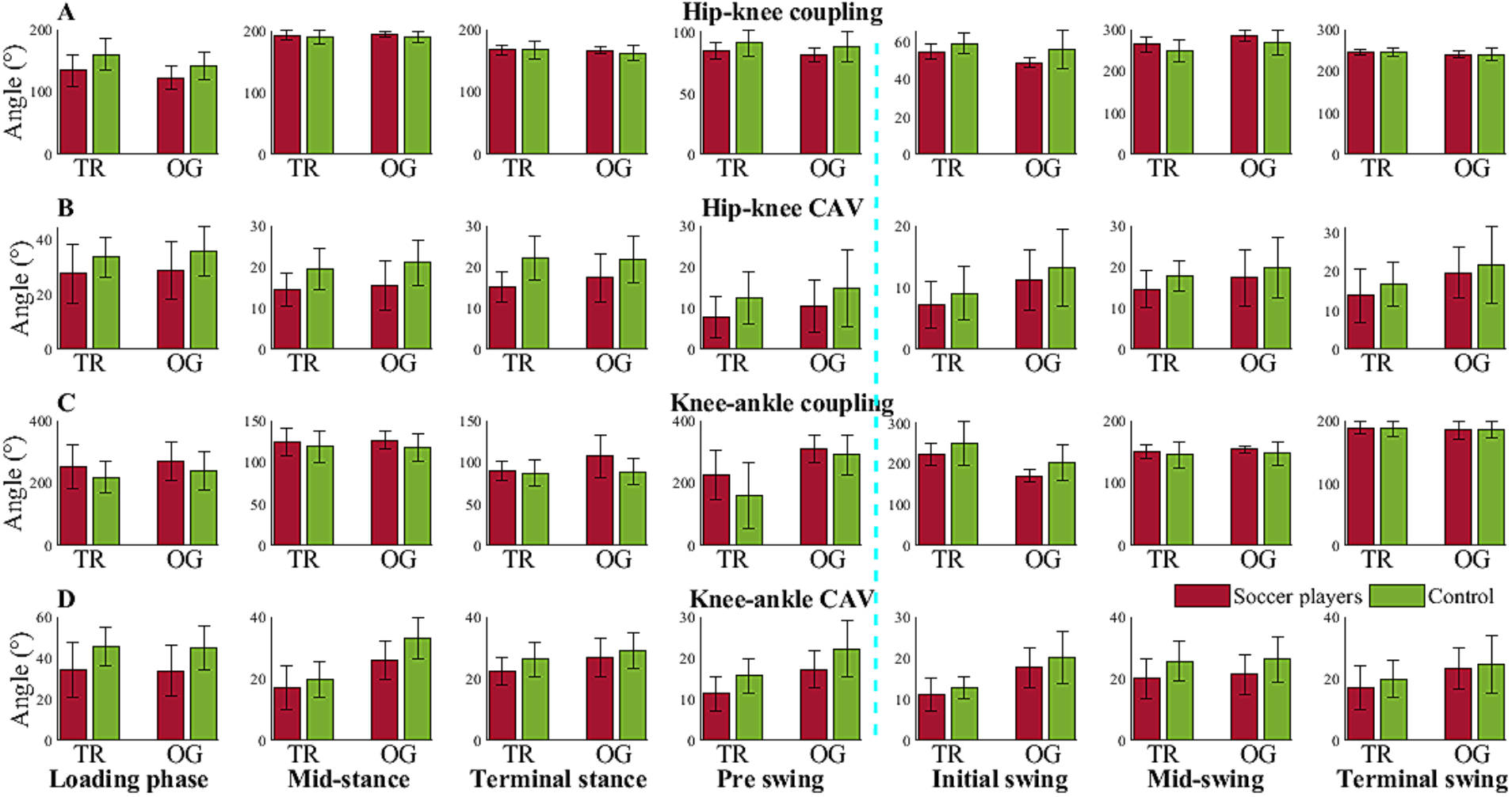
Coupling angle and CAV of soccer players and control group during walking on treadmill or overground. Rows A and B: hip-knee coupling angle and CAV, respectively. Rows C and D: knee-ankle coupling angle and CAV, respectively. Each row is divided into seven sub-phases of gait cycle. Blue dash line indicates toe-off and separates the stance phase (left side) and swing phase (right side). Each subplot shows mean and SD of coupling/CAV of athletes and non-athlete during treadmill (TR) or overground (OG) walking.

## 4. Discussion

We used a vector coding technique to compare the coordination of hip, knee and ankle joint angles from the right leg in the sagittal plan between soccer players and controls when walking overground or on a treadmill. We observed differences in CAV between groups in different phases of the gait cycle. Soccer players showed decreased hip-knee CAV during the mid-stance and terminal-stance phases and decreased knee-ankle CAV during the pre-swing phase compared to the control group. The mean coupling angle was used to distinguish between four coordination patterns: in-phase, anti-phase, proximal phase and distal phase. This revealed that the control group more often showed a distal-phase (ankle) coordination pattern for the knee-ankle pair compared to soccer players. These findings show that the vector coding technique can detect differences in joint coordination and variability during normal gait between soccer players and controls.

Soccer players had lower hip-knee CAV during middle stance and lower knee-ankle CAV during the pre-swing phase compare to the control group. Recently, the functional role of variability in system dynamics has been discussed and that the traditional view of disordered movement as showing more variability has been questioned (Hamill, van Emmerik, Heiderscheit, & Li, 1999). They suggested a functional role for variability in lower extremity segment coupling in which symptomatic individuals applied joint actions within a very narrow range that led to less variability compared to healthy subjects. This raises the question why variability is lower in both pathological condition as shown in previous studies and in trained healthy individuals (soccer players) in our study. A lower CAV in symptomatic individuals may help them to minimize the pain during movement, but both groups in our study were functionally able to perform walking without pathological symptoms. While we are not aware of any study that has investigated CAV of the lower extremity in the gait pattern of soccer players, some studies have assessed the variability of kinematic variables during sports movement. For example, the least, intermediate and the most skilled triple jumpers exhibited U-shape curve in coordination variability during hop-step phase, as skill increased. The authors explained that high coordination variability may not be beneficial in the least skilled jumpers and suggested that variability decreases when jumpers are able to demonstrate more consistent or regulated performance (Wilson et al., 2008). Overall, less experienced individuals show more variability on a given task (Jarvis, Smith, & Kulig, 2014), the extend of which decreases as they learn and approach expert performances. Therefore, decreased CAV in soccer players may reflect adaptation towards optimal variability (Stergiou, Harbourne, & Cavanaugh, 2006) that improves gait stability (Hyun G Kang & Jonathan B Dingwell, 2008; Hyun Gu Kang & Jonathan B Dingwell, 2008; Owings & Grabiner, 2004) as a result of soccer training. However, we should be aware that the most experienced individuals may exhibit a degree of variability that allows them flexibility in dealing with perturbations and controlling balance, timing, and any other applicable factor (Wilson et al., 2008).

Our findings also showed that soccer players showed less ankle coordination pattern (distal phase) compared to the control group and that this difference was most evident in the stance phase (Fig. 4). A knee flexion is essential for energy absorption and is relevant to joint degeneration during leading phase of gait (Childs, Sparto, Fitzgerald, Bizzini, & Irrgang, 2004). In addition, the ankle joint moments revealed that the greatest demands on the controlling muscles occurred during stance phase of gait (Hunt, Smith, & Torode, 2001). Indeed, the ankle joint muscles have a crucial function in stabilizing the foot when weight transfers onto the toes (Hunt & Smith, 2004). Therefore, the ability to coordinate the movement of the knee and ankle joints is important during the stance phase of gait. Soccer players should properly position their feet during soccer training/competition (Hawrylak, Brzeźna, & Chromik, 2021), therefore, gait motor control of soccer players may be unconsciously trained for a better foot position. According to the present results, controls tried to preserve greater ankle coordination pattern to transfer the body weight during stance phase. However, soccer players may show good coordination (anti-phase coupling) in knee-ankle coupling, where knee flexion versus ankle extension leads to better load absorption and weight control during stance phase (Fig. 4F).

It should be noted that this study was conducted with a small sample size and assessed many outcome variables. We did use the Benjamini-Hochberg procedure to control the false discovery rate, but our findings should be replicated in future studies involving a larger sample to confirm that the stance phase is important part of gait cycle that is affected by long-term training (Kiriyama, Warabi, Kato, Yoshida, & Kokayashi, 2005). In addition, this study conducted using only male participants, while previous studies demonstrated gender-related differences in lower limb variability (Barrett, Noordegraaf, & Morrison, 2008; Kerrigan, Todd, & Della Croce, 1998). Females showed less variability in ankle transfer plan rotations compared to males at different speeds of treadmill running, and females also had greater hip flexion or knee extension before initial contact. Lastly, we did not assess kinetics and neuromuscular control mechanisms, which may reveal further insights into the mechanism underlying the observed difference in joint coordination. In addition to lower extremity kinematics, future studies should attend to other aspects of gait motor control, kinetics and electromyography, to reveal more accurate information of gait motor control. Therefore, extrapolating the conclusions from our study to other gender group (females) and larger group (participants number) must be done with caution.

## 5. Conclusions

In summary, we found that lower extremity joint variability and coordination during walking differs between soccer players and controls and may thus vary even among healthy young adults. This differences in coordination could be related to the physical fitness background of the participants. Soccer players showed lower hip-knee CAV during mid-stance and terminal stance phases and lower knee-ankle CAV during the pre-swing phase of gait, and also applied less ankle coordination pattern compared to the control group. We hypothesized that long-term soccer training could be one of the reasons for these differences, which may reflect adaptations to exercise training by reducing variability during the stance phase of gait. Future efforts should attempt to evaluate this hypothesis by examining the changes in joint coordination of the lower extremity before and after an exercise training intervention.

## Acknowledgments

We would like to thank the volunteer participants for their participation in the experiments and the Otech_motionlab group for their cooperation. Tjeerd Boonstra has received funding from the European Union’s Horizon 2020 research and innovation programme under the Marie Sklodowska-Curie grant agreement No 895914.

## Author contribution

Z.F.M, M.Y, S.E.H, and I.E.P.A conceived the study. M.Y. performed the experiments. M.R, T.W.B, and K.M carried out the analyses. M.Y. and T.W.B wrote the main text. All authors reviewed and edited the manuscript.

