## Supplementary Material for "The coordination of hip, knee and ankle joint angles during gait in soccer players and controls"

Table S1. Statistical results of hip-knee and knee-ankle coupling and CAV of gait cycle during walking with walking speed as covariate variable.

| Variable / Condition | Hip-knee coupling | | | Knee-ankle coupling | | | Hip-knee CAV | | | Knee-ankle CAV | | |
|---|---|---|---|---|---|---|---|---|---|---|---|---|
| | F | P | $P_{adj}$ | F | P | $P_{adj}$ | F | P | $P_{adj}$ | F | P | $P_{adj}$ |
| Loading response (1-10%) | | | | | | | | | | | | |
| Group | 5.15 | 0.03 | 0.25 | 0.69 | 0.41 | 0.87 | 4.81 | 0.03 | 0.25 | 8.18 | 0.01 | 0.11 |
| Surface | 3.81 | 0.06 | 0.31 | 2.15 | 0.15 | 0.48 | 0.91 | 0.34 | 0.85 | 1.33 | 0.26 | 0.67 |
| In G * S | 0.27 | 0.61 | 0.92 | 0.002 | 0.97 | 0.98 | 0.06 | 0.81 | 0.92 | 0.001 | 0.97 | 0.98 |
| Mid-stance (11-30%) | | | | | | | | | | | | |
| Group | 2.63 | 0.11 | 0.38 | 3.23 | 0.08 | 0.32 | **14.70** | **<0.001** | **0.008** | 7.93 | 0.01 | 0.11 |
| Surface | 0.52 | 0.47 | 0.89 | 0.11 | 0.74 | 0.92 | 0.03 | 0.85 | 0.93 | 1.89 | 0.18 | 0.54 |
| In G * S | 0.16 | 0.69 | 0.92 | 0.08 | 0.78 | 0.92 | 0.01 | 0.92 | 0.96 | 0.21 | 0.65 | 0.92 |
| Terminal stance (31-50%) | | | | | | | | | | | | |
| Group | 0.83 | 0.37 | 0.87 | 3.95 | 0.05 | 0.31 | **18.24** | **<0.001** | **0.003** | 5.26 | 0.03 | 0.25 |
| Surface | 0.06 | 0.81 | 0.92 | 0.45 | 0.51 | 0.92 | 0.18 | 0.67 | 0.92 | 0.12 | 0.74 | 0.92 |
| In G * S | 0.40 | 0.53 | 0.92 | 2.94 | 0.09 | 0.35 | 0.85 | 0.36 | 0.87 | 0.41 | 0.52 | 0.92 |
| Pre swing (51-60%) | | | | | | | | | | | | |
| Group | 3.57 | 0.06 | 0.31 | 0.55 | 0.46 | 0.89 | 3.65 | 0.06 | 0.31 | **12.18** | **0.001** | **0.02** |
| Surface | 0.76 | 0.39 | 0.87 | 0.003 | 0.96 | 0.98 | 2.20 | 0.14 | 0.47 | 3.53 | 0.07 | 0.32 |
| In G * S | 0.02 | 0.90 | 0.95 | 1.41 | 0.24 | 0.67 | 0.01 | 0.91 | 0.95 | 0.06 | 0.81 | 0.92 |
| Initial swing (61-73%) | | | | | | | | | | | | |
| Group | 5.74 | 0.02 | 0.24 | 3.48 | 0.07 | 0.32 | 3.63 | 0.06 | 0.31 | 5.31 | 0.03 | 0.25 |
| Surface | 0.13 | 0.72 | 0.92 | 0.31 | 0.58 | 0.92 | 0.18 | 0.67 | 0.92 | 1.71 | 0.20 | 0.58 |
| In G * S | 0.83 | 0.37 | 0.87 | 0.03 | 0.86 | 0.93 | 0.00 | 0.99 | 0.99 | 0.12 | 0.73 | 0.92 |
| Mid-swing (74-87%) | | | | | | | | | | | | |
| Group | 3.10 | 0.08 | 0.32 | 0.71 | 0.40 | 0.87 | 3.11 | 0.08 | 0.32 | 5.10 | 0.03 | 0.25 |
| Surface | 0.16 | 0.69 | 0.92 | 0.71 | 0.41 | 0.87 | 0.63 | 0.43 | 0.89 | 1.82 | 0.18 | 0.54 |
| In G * S | 0.10 | 0.76 | 0.92 | 0.07 | 0.79 | 0.92 | 0.06 | 0.81 | 0.92 | 0.03 | 0.86 | 0.93 |
| Terminal swing (88-100%) | | | | | | | | | | | | |
| Group | 0.15 | 0.70 | 0.92 | 2.85 | 0.10 | 0.37 | 3.34 | 0.07 | 0.32 | 2.17 | 0.15 | 0.48 |
| Surface | 0.07 | 0.80 | 0.92 | 7.07 | 0.01 | 0.14 | 0.02 | 0.89 | 0.95 | 0.20 | 0.66 | 0.92 |
| In G * S | 0.05 | 0.82 | 0.92 | 0.20 | 0.66 | 0.92 | 0.06 | 0.81 | 0.92 | 0.21 | 0.65 | 0.92 |

$P_{adj}$ = P-values are adjusted using the Benjamini-Hochberg procedure, CAV = Coupling angle variability, In G*S = Interaction effect of group * surface. Significant effects (P<0.05) are displayed in bold.

Table S2. Descriptive statistics of coordination frequency for four phases

| Condition / Phase | Athlete | | Non-athlete | | Treadmill | | Overground | |
|---|---|---|---|---|---|---|---|---|
| | mean | SD | mean | SD | mean | SD | mean | SD |
| Anti-phase H-K | 18.21 | 4.01 | 17.19 | 5.72 | 16.40 | 5.43 | 18.93 | 4.20 |
| In-phase H-K | 28.96 | 6.81 | 23.28 | 10.22 | 25.87 | 9.67 | 26.00 | 8.83 |
| Hip phase H-K | 23.21 | 10.10 | 31.75 | 10.79 | 29.17 | 12.10 | 26.37 | 10.32 |
| Knee phase H-K | 29.61 | 6.58 | 27.78 | 5.53 | 28.57 | 5.90 | 28.70 | 6.32 |
| Anti-phase K-A | 31.04 | 7.03 | 23.31 | 10.14 | 24.07 | 9.67 | 29.77 | 8.74 |
| In-phase K-A | 12.46 | 2.63 | 12.72 | 2.47 | 13.57 | 2.69 | 11.63 | 1.96 |
| Knee phase K-A | 45.07 | 8.02 | 43.25 | 5.55 | 44.80 | 6.97 | 43.40 | 6.71 |
| Ankle phase K-A | 11.43 | 5.44 | 20.72 | 9.26 | 17.57 | 9.59 | 15.20 | 8.29 |

H-K = Hip-Knee, K-A = Knee-Ankle.

Table S3. Statistical results of coordination frequency of hip-knee and knee-ankle couplings with walking speed as covariate variable.

| Condition / Variable | Group | | | Surface | | | Group * Surface | | |
|---|---|---|---|---|---|---|---|---|---|
| | F | P | $P_{adj}$ | F | P | $P_{adj}$ | F | P | $P_{adj}$ |
| Anti-phase Hip-Knee | 0.07 | 0.80 | 0.92 | 1.38 | 0.25 | 0.67 | 0.34 | 0.57 | 0.92 |
| In-phase Hip-Knee | 4.65 | 0.04 | 0.27 | 0.03 | 0.85 | 0.93 | 2.64 | 0.11 | 0.38 |
| Hip phase Hip-Knee | 4.61 | 0.04 | 0.27 | 1.36 | 0.25 | 0.67 | 0.68 | 0.41 | 0.87 |
| Knee phase Hip-Knee | 0.46 | 0.50 | 0.92 | 0.61 | 0.44 | 0.89 | 0.27 | 0.60 | 0.92 |
| Anti-phase Knee-Ankle | 7.81 | 0.01 | 0.11 | 0.12 | 0.73 | 0.92 | 0.46 | 0.50 | 0.92 |
| In-phase Knee-Ankle | 0.55 | 0.46 | 0.89 | 0.56 | 0.46 | 0.89 | 0.27 | 0.61 | 0.92 |
| Knee phase Knee-Ankle | 0.35 | 0.56 | 0.92 | 1.28 | 0.26 | 0.67 | 0.14 | 0.71 | 0.92 |
| Ankle phase Knee-Ankle | **14.43** | **<0.001** | **0.008** | 0.17 | 0.68 | 0.92 | 0.30 | 0.59 | 0.92 |

$P_{adj}$ = adjusted. * P-values are adjusted using the Benjamini-Hochberg procedure. Significant effects ($P<0.05$) are displayed in bold.

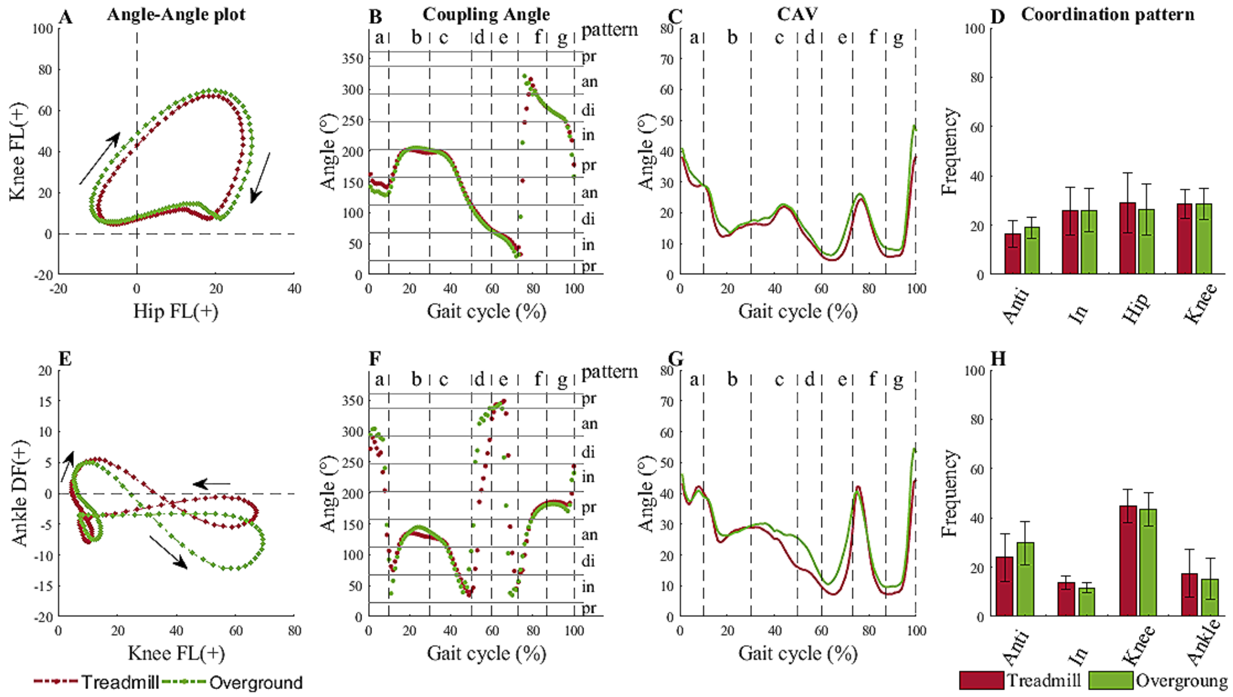


Fig. S2. The mean of hip-knee and knee-ankle coordination in sagittal plan for treadmill and overground players and controls. First column from left shows the angle-angle plots for hip-knee and knee-ankle coordination with proximal joint on the horizontal axis. The second column shows hip-knee and knee-ankle coupling angles. Note that the vector coding procedure yields coupling angle values between 0° - 360°. Vertical dash lines divided gait cycle to seven sub-phases and horizontal grey solid lines shows the associated coordination patterns (anti-phase, in-phase, proximal segment phase, distal segment phase). The third column shows coupling angle variability which is divided in gait cycle sub-phases (vertical dash lines). The fourth column shows the average frequency (%) of each coordination patterns and the error bars show the standard deviation between participants. Anti = anti-phase, In = in-phase, a = loading response, b = mid-stance, c = terminal stance, d = pre-swing, e = initial swing, f = mid-swing, g = terminal swing. * = significant difference ($P<0.05$) in coordination pattern. The cyan bands show the sub-phase with a significant main effect on CAV.
